# High angular resolution diffusion MRI reveals conserved and deviant programs in the paths that guide human cortical circuitry

**DOI:** 10.1101/576967

**Authors:** Christine J. Charvet, Avilash Das, Jae W. Song, Deselyn J. Tindal-Burgess, Priya Kabaria, Guangping Dai, Tara Kane, Emi Takahashi

**Author notes:** Corresponding authors: Christine Charvet, Ph.D., Department of Psychology, Delaware State University, Dover, DE, 19901, USA, Emi Takahashi, Ph.D., Division of Newborn Medicine, Boston Children’s Hospital, Harvard Medical School, 401 Park Dr., Boston, MA 02215, USA.

## Abstract

Diffusion MR tractography represents a novel opportunity to investigate conserved and deviant developmental programs between humans and other species such as mice. To that end, we acquired high angular resolution diffusion MR scans of mice (embryonic day [E] 10.5 to post-natal week [PW] 4) and human brains (gestational week [GW] 17 to 30) at successive stages of fetal development to investigate potential evolutionary changes in radial organization and emerging pathways between humans and mice. We compare radial glial development as well as commissural development (e.g., corpus callosum), primarily because our findings can be integrated with previous work. We also compare corpus callosal growth trajectories across primates (i.e., humans, rhesus macaques) and rodents (i.e., mice). One major finding is that the developing cortex of humans is predominated by pathways likely associated with a radial glial organization at GW 17-20, which is not as evident in age-matched mice (E 16.5, 17.5). Another finding is that, early in development, the corpus callosum follows a similar developmental timetable in primates (i.e., macaques, humans) as in mice. However, the corpus callosum grows for an extended period of time in primates compared with rodents. Taken together, these findings highlight deviant developmental programs underlying the emergence of cortical pathways in the human brain.

## Introduction

Diffusion MR imaging is a high-throughput and non-invasive method that can be used to investigate neural pathways in adulthood, as well as their development (Mori and Zhang, 2006; Qui et al., 2015). Questions focused on the development and evolution of connectivity patterns and cell migration have traditionally been addressed with tract-tracers or carbocyanine tracers. Although these methods have revealed key developmental processes in the brain, they are time-consuming, invasive, and suffer from a number of technical limitations (Nudo and Masterton, 1989, 1990; Chen et al., 2006; Heilingoetter and Jensen, 2016). Moreover, carbocyanine tracers (e.g., DII) are one of the few methods available for study of migration and establishment of pathways in humans (Burkhalter et al., 1993; Konstantinidou et al., 1995; Hevner, 2000; Tardif and Clarke, 2001). As a result, we have been able to only glimpse at the developmental processes dictating evolutionary changes in patterns of brain connections in the human lineage.

Diffusion imaging has revealed important developmental processes in humans such as radial glial pathways in the developing cortex (Takahashi et al., 2012; Kolasinski et al., 2013; Xu et al., 2014; Miyazaki et al., 2016; Vasung et al., 2017; Das and Takahashi, 2018), tangential migratory routes (Takahashi et al., 2012; Kolasinski et al., 2013; Miyazaki et al., 2016; Wilkinson et al., 2017), and emerging axonal pathways coursing through the white matter of the brain (e.g. Vasung et al., 2010; Takahashi et al., 2012). We here focus on the developmental timeline of radial glia and commissural fibers in humans and mice to identify conserved and deviant developmental programs giving rise to evolutionary changes in the structure and connectivity patterns in the human brain. Among the commissural fibers available for study, we focus especially on the developmental timeline of the corpus callosum in humans and mice because the corpus callosum is clearly homologous across rodents and primates, which is not true of many other cortical association pathways (e.g., uncinate fasciculus, arcuate fasciculus; Schmahmann et al., 2007; Charvet, Šimić, et al., 2017). The corpus callosum is a white matter fiber pathway integral for inter-hemispheric cortical communication, which evolved with the emergence of placental mammals, and is disproportionately expanded in primates compared with many other eutherian mammals (Katz et al., 1983; Aboitiz and Montiel, 2003; Bloom and Hynd, 2005; Manger et al., 2010; Caminiti et al., 2013). Diffusion MR tractography provides a three-dimensional perspective with which to compare development of radial fibers and emerging axonal pathways across the entire cortex of mammalian species.

A complex process orchestrates cell production in the human brain. In humans, neurogenesis across the six layered cortex (called the isocortex) is thought to begin at around embryonic day E 30 (Bystron et al., 2006), and ceases around birth (270 days after gestation; Malik et al., 2013; Zecevic et al., 2005; Charvet, Hof et al., 2017), whereas isocortical neurogenesis in mice begins on E10 to last only about 8 days (Caviness et al., 2003; 2009). As cells exit the proliferative zone to become neurons, they migrate towards the cortical plate (CP) along radial glia (Varon and Somjen, 1979; Rakic, 1971, 1972, 1974, 2002, 2003; Patel et al., 1994, Noctor et al., 2001; Rash et al., 2001; deAzevedo et al., 2003; Getz et al., 2014; Nowakowski et al., 2016). The first neurons to extend axons are called pioneers. Similar to radial glial cells, which guide the migration of newly born neurons, pioneer axons act as scaffolds guiding the extension of other axons (Bently and Keshishian, 1982; Supér et al., 1998; Nishikimi et al., 2013). Pioneer axons guide the development of cortical neurons projecting subcortically and cortically, including neurons projecting to the contralateral hemisphere (McConnell et al., 1989; De Carlos and O’Leary, 1992; Koester and O’Leary, 1994; Rash and Richards, 2001; Paul et al., 2007). As more and more cells exit the cell cycle, the progenitor pool, and the radial organization of the developing human fetal cortex wane, and the white matter of the developing cortex becomes increasingly replaced with long-range projecting pathways as assessed from diffusion MR imaging (Takahashi et al., 2012). The aim of the present study is to compare the timetable of “scaffolds” such as radial glial fibers and pioneer axons between species. Although “pioneer” might imply a guidance function, we define pioneer exclusively temporally, and we focus on the temporal sequence of fibers emerging across the hemisphere of the developing cortex. Another aim of the present study is to compare the timing of cortical association pathway maturation (i.e., corpus callosum) between human and mice to investigate evolutionary changes in developmental processes giving rise to variation in cortico-cortical association pathways between species.

It is well known that primate isocortical neurogenesis is unusual compared with rodents (Rakic, 1995; Zecevic et al., 2005; Kriegstein et al., 2006; Martínez-Cerdeño et al., 2006). After controlling for differences in developmental schedules, isocortical neuronal production extends for an unusually long time in humans and macaques compared with mice (Clancy et al., 2001; Workman et al., 2013; Cahalane et al., 2014; Charvet, Hof, et al., 2017). The extension in the duration of isocortical neurogenesis leads to an amplification of upper layer neurons (i.e., layer II-IV), and a concomitant expansion of long-range cortico-cortical association pathways in adult primates (Charvet et al., 2015, 2019; Charvet, Hof, et al., 2017; Krienen et al., 2016). Given these data, it may be expected that there are differences in radial glial organization between humans and mice during neurogenesis because an extension in the duration of cortical neurogenesis should entail an extended period in which fibers are radially organized to allow newly born neurons to migrate from the progenitor pool to the CP. Extending the duration of neurogenesis might also lead to major changes in cortical circuitry by potentially altering the timing of initial axon extension of pioneer neurons, and/or extending the duration over which cortical association fibers grow. Many of the collosally-projecting neurons are situated in upper layer (i.e., layer II-IIV) neurons (Isseroff et al. 1984; Meissirel et al. 1991; Greig et al., 2013), which have undergone a dramatic expansion in primates (Charvet et al., 2017; 2019), which is concomitant with increased expression of select genes encoding ion channels, neurofilament and synaptic proteins in upper layers (Zeng et al., 2012; Krienen et al., 2016). In addition, callosally-projecting neurons express a suite of genes, some of which are differentially expressed in macaques compared with mice (Fame et al., 2011, Fame, Dehay, et al., 2016). These studies highlight major modifications in callosal organization between primates and rodents. Thus, we can use diffusion MR tractography to investigate deviations in developmental processes giving rise to the evolution of cortical association pathways in the human lineage.

Considerable work has been conducted on radial glial development, pioneer, as well as callosal development in model organisms such as mice, rats, monkeys, and cats (Wise and Jones, 1976; Marin-Padilla, 1978; Innocenti, 1981; O’Leary et al., 1981; Ivy and Killackey, 1982; Silver et al., 1982; Luskin and Shatz, 1985; Berbel and Innocenti, 1988; Mrzljak et al., 1988; Voigt, 1989; Mrzljak et al., 1990; Lent et al., 1990; LaMantia and Rakic, 1990; Aggoun-Zouaoui and Innocenti, 1994; Tessier-Lavigne and Goodman, 1996; Richards et al., 1997; Shu and Richards, 2001; Richards, 2002, 2012; Unni et al., 2012; Fame, MacDonald, et al., 2016; Gobius et al., 2016; Niquille et al., 2019), as well as humans (Rakic and Yakovlev, 1968; Kostovic and Krmpotic, 1976; Kostovic and Rakic, 1990; Marin-Padilla, 1992; Kier and Truwit, 1996; deAzevedo et al. 1997; Rados et al. 2006; Ren et al., 2006; Huang et al. 2006; Huang et al. 2009). Yet, we still know very little about the basic developmental timeline of radial glia, pioneer development, and cortical association pathways in humans, and which developmental processes are conserved or variant between humans and other species. Understanding basic developmental differences between humans and model organisms is an essential enterprise in order to relate findings from model organisms to humans, and to identify developmental mechanisms that underlie the neural specializations responsible for sensory and cognitive specializations leading to the emergence of the human brain (Clancy et al., 2001; Ward and Vallender, 2012; Perlman, 2016; Charvet, Šimić, et al., 2017).

Among the various diffusion MR methods amenable to reconstruct pathways in three dimensions, high-angular resolution diffusion imaging (HARDI) tractography enables the identification of complex crossing tissue coherence in the brain (Tuch et al., 2003), as well as immature brains (e.g. Takahashi et al., 2010, 2011, 2012). It is typically more challenging to identify cell extensions whether they are from radial glia or from bundles of axons because there is a surplus of unmyelinated fibers during fetal brain development. We here use HARDI to track the development of fibers because it is theoretically superior to diffusion tensor imaging (DTI), and permits the study of pathways guiding the development of neural pathways (e.g., radial glia, pioneer axons) in the human brain.

## Materials and Methods

### Sample acquisition for diffusion MR tractography

Five intact mouse brains at E 10.5, 14.5, 16.5, 17.5, and 4 postnatal weeks (PW) were obtained from ongoing developmental or genetic research projects at Boston Children’s Hospital (BCH) and Massachusetts General Hospital (MGH) under approved protocols at the hospitals. Mice were deeply anesthetized (Xylanzine/Ketamine 100 mg/ml, 60 mg/Kg) and either sacrificed by transcardial perfusion with 0.9% saline followed by 4% paraformaldehyde (>post-natal day 5), or decapitated and immersion fixed (<post-natal day 3). Fetal brains were immersion fixed after we removal of the skull. Postnatal animals were perfused with 4% paraformaldehyde followed by immersion fixation for at least 2 weeks in 4% PFA.

A total of five brains at GW 17, 20, 21, 22, and 30 were imaged for diffusion MR tractography. Samples from GW17-22 were scanned *ex-vivo* whereas the GW30 individual was scanned *in vivo*. Postmortem brains were obtained from the Department of Pathology, Brigham and Women’s Hospital (BWH; Boston, MA, USA) and the Allen Institute Brain Bank (AIBB; Seattle, WA, USA) with full parental consent. The known primary causes of death were complications from prematurity. The brains were grossly normal, and standard autopsy examinations of all brains undergoing postmortem HARDI revealed no or minimal pathologic abnormalities at the macroscopic level. All brains utilized for this study were analyzed under institutional review board approved protocols.

### Tissue preparation for HARDI

At the time of autopsy, all brains were immersion fixed and the brains from the BWH were stored in 4% paraformaldehyde, and the brains from the AIBB were stored in 4% periodate-lysine-paraformaldehyde (PLP). During MR image acquisition, the brains from the Brigham and Women’s Hospital were placed in Fomblin solution (Ausimont, Thorofare, NJ; e.g. Takahashi et al., 2012) and the brains from the AIBB were placed in 4% PLP. These different kinds of solutions in which the brains from different institutes were placed tend to change the background contrast (i.e. a dark background outside of the brain using Fomblin, and bright background using PLP), but they do not specifically change diffusion properties (e.g. fractional anisotropy: FA and apparent diffusion coherence: ADC) within the brain.

### Diffusion MRI procedures

Different scanner systems were used to accommodate brains of various sizes (same as in e.g. Takahashi et al., 2012; Xu et al., 2014; Miyazaki et al., 2015; Wilkinson et al., 2017). Magnetic resonance (MR) coils that best fit each brain sample were used to ensure optimal imaging. Mouse specimens were imaged with a 9.4T Bruker Biospec MR system. The postmortem brain specimens from BWH were imaged with a 4.7T Bruker Biospec MR system (2 fetal specimens [GW 17, 20]), and specimens from the AIBB were imaged with a 3T Siemens MR system (2 fetal specimens [GW 21, 22]) at the A. A. Martinos Center, MGH, Boston, MA, USA. The 3T system was used to accommodate the AIBB brains that were in cranium and did not fit in the 4.7T bore. In order to obtain the best signal-to-noise ratio and highest spatial resolution, we used custom-made MR coils with one channel on the 4.7T and 3T systems (Takahashi et al. 2012; Kolasinski et al. 2013; Xu et al. 2014).

To supplement the small sample size, we used brains from two different sources. Because brains from one of the sources (GW 21 and GW 22) had skulls, a different scanner system and scan parameters had to be used. Only two ex vivo specimens at GW 21 and GW 22 were scanned at 3T, because the samples with skulls do not fit into the 4.7T scanner. For the *in vivo* subject, the 4.7T small-bore scanner cannot be used, so a 3T scanner was used. At a 3T scanner, the use of high b values such as 8000s/mm^2^ is problematic because it causes artifacts in that the tractography appears inconsistent with previous findings from tract-tracers. We reason that the issue of different acquisition systems is not significant for tractography because both systems yielded similar patterns.

For the BWH brains and mouse specimens, a three-dimensional (3-D) diffusion-weighted spin-echo echo-planar imaging (SE-EPI) sequence was used with a repetition time/echo time (TR/TE) of 1000/40 ms, with an imaging matrix of 112 × 112 × 112 pixels. Sixty diffusion-weighted measurements (with the strength of the diffusion weighting, b = 8000 s/mm^2^) and one non-diffusion-weighted measurement (no diffusion weighting or b = 0 s/mm^2^) were acquired with δ = 12.0 ms and Δ = 24.2 ms. The spatial resolution was 440 × 500 × 500 µm. For the brains from the AIBB, diffusion-weighted data were acquired over two averages using a steady-state free-precession sequence with TR/TE = 24.82/18.76 ms, α = 60°, and the spatial resolution was 400 × 400 × 400 µm. Diffusion weighting was isotropically distributed along 44 directions (b = 730 s/mm^2^) with 4 b = 0 images, using a custom-built, single channel, 2-turn solenoid coil (6.86 cm inner diameter, 11.68 cm length). The solenoid design minimizes the distance between the copper turns of the coil and the sample, yielding higher signal-to-noise ratio with each turn, and improved signal uniformity compared with a phased array receive coil design. We determined the highest spatial resolution for each brain specimen with an acceptable signal-to-noise ratio of more than 130 for diffusion MR tractography.

The in vivo subject (GW 30) from BCH was imaged on a 3T Siemens MR system at Boston Children’s Hospital, Boston, MA, USA when the individual was asleep and not anaesthetised. The diffusion pulse sequence used was a diffusion-weighted SE-EPI sequence, TR/TE 8320/88 ms, with an imaging matrix of 128 × 128 × 64 pixels. The spatial resolution was 2 × 2 × 2 mm. Thirty diffusion-weighted measurements (b = 1000 s/mm2) and 5 b = 0 images were acquired with δ = 40 ms and Δ = 68 ms.

### Reconstruction and identification of diffusion MR tractography

We reconstructed white matter tracts in each brain with the Diffusion Toolkit and TrackVis (http://trackvis.org). We used a FACT algorithm for diffusion MR tractography (Mori et al. 1999), as in previous publications (for humans, Takahashi et al, 2010, 2011, 2012; Schmahmann et al, 2007; D’Arceuil et al, 2008; Das and Takahashi 2018; for mice, Rosen et al., 2013; Lodato et al., 2014; Fame, Dehay, et al., 2016; Kanamaru et al., 2017), which connects fibers using a local maximum or maxima. The FACT algorithm is limited in its ability to resolve crossing pathways when used with the traditional DTI technique because one simply connects the direction of the principal eigenvector on a tensor to produce the DTI tractography pathways. This feature is a recognized limitation of DTI (Mori et al., 1999). Hence, in the current study, we used HARDI, which can theoretically detect multiple local maxima on an orientation distribution function (ODF). Using each local maxima on an ODF, we applied the streamline algorithm to initiate and continue tractography (Tuch et al., 2003), thus enabling us to identify crossing pathways within a voxel.

Diffusion MR trajectories were propagated by consistently pursuing the orientation vector of least curvature. We terminated tracking when the angle between two consecutive orientation vectors was greater than the given threshold (40°), or when the fibers extended outside of the brain surface. To study fibers in contact with the medial walls of the developing cortex, multiple sagittal slice filters of varying thickness with “either end” restrictions were used. Thickness of slice filters and placement was altered depending on the fiber population of interest. We also used whole brain MR tractography with minimum length threshold set to 0.8mm in the mouse brain, and we placed ROIs to highlight fibers of interest. For visualization purposes, the color-coding of tractography connections was based on a standard red-green-blue (RGB) code using directions of the middle segment of each fiber, and the color coding in humans was based on an RGB code. In some cases, this RGB code relies on the direction of every fiber segment. In other cases, the RGB code is applies to the average direction of fibers.

### Variation in corpus callosum area between primates and rodents

It has been previously reported that the corpus callosum is expanded in primates compared with other mammals (Manger et al., 2010). We aimed to confirm that the corpus callosum is indeed expanded in primates compared with rodents. To that end, we use structural MRI and atlases to measure the corpus callosum area from mid-sagittal sections in small-brained monkeys and large-brained rodents in order to control for variation in overall brain size. We supplement these data with previously published data (Charvet et al., 2019; Manger et al., 2010; Yuasa and Kohsaka, 2010; Schilling et al., 2017; Charvet et al., 2015). We use phylogeny-generalized-least-squares statistics (PGLS) to obtain phylogenetically-controlled slopes of the logged values of the area of corpus callosum versus the logged values of brain size (Pagel, 1999; Freckleton et al., 2000). We test whether taxonomy (i.e, primate versus rodent) accounts for a significant percentage of the variation in corpus callosum area. These phylogenetically-controlled generalized linear models were performed via restricted estimated maximum likelihood with the software program R, and the library package ape (ML; Pagel, 1999; Freckleton et al., 2000). The phylogeny, which includes branch lengths for these species, was taken from Bininda-Emonds et al., 2007.

After controlling for variation in brain size, primates possess a larger corpus callosum compared with rodents (**Supplementary Fig. 1**). A PGLS model with brain size and taxonomy are both significant independent variables accounting for 98.2% of the variance in corpus callosum area (F= 348; R^2^=0.98.1; p=1.137e-10; λ=0) and an AIC score of −25.51. A PGLS model that uses brain size alone to predict corpus callosum area only accounts for 96.8% of the variance (F= 393.3; p= 1.539e-10, λ=0), with an AIC score model of −18.50. The addition of taxonomy as an independent variable in the PGLS model results in a lower AIC score than the model without taxonomy as an independent variable. These findings confirm that primates possess a larger corpus callosum than rodents when controlled for brain size.

**Figure 1.**
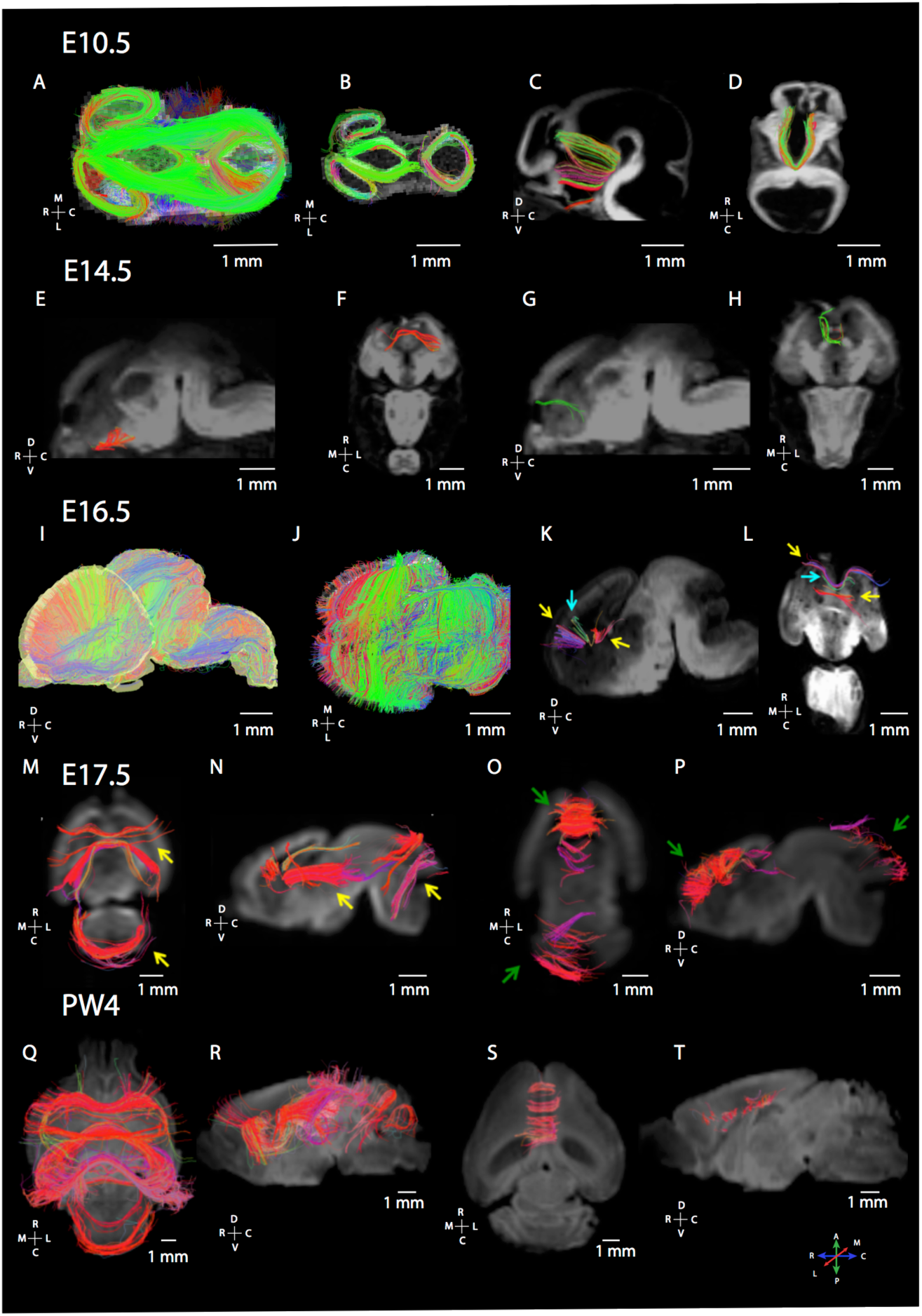
Maturation of commissural fibers as assessed from diffusion MR tractography in mice at various age of development (E10.5, E14.5, E16.5, E17.5, post-natal week 4: PW4). (A) At E10.5 (A-D), dorsal view of brain fibers (A-B) show that many of the pathways course across the anterior to posterior direction in the dorsal telencephalon as well as in the diencephalon. We identify pathways coursing through the midline in the ventral telencephalon (C, D). (B) At E14.5 (E-H), we observe callosal pathways as well as putative pioneer axons. (C) At E16.5 (I-L), we observe pathways coursing across the medial to lateral axes through the developing cortex (I-J). At these ages, we also identify callosal and putative pioneer pathways. Yellow arrows in 16.5E denote putative pioneer axons, while blue arrows denote callosal pathways. At E17.5 (M-P), horizontal (M, O), and sagittal views (N, P) highlight callosal and putative pioneer pathways coursing through the dorsal midline. At post-natal week 4 (Q-T), we observe the corpus callosum as well as the hippocampal commissure coursing across the medial to lateral direction within the caudal cortex. Yellow arrows show the development of fibers in the anterior and posterior brain regions, while green arrows highlight the development of short fibers in the anterior and posterior brain regions. The direction of fibers is color-coded with a map of these color-codes located in the lower left panel. An orientation map is adjacent to each examined plane of section. The following abbreviations are used: R: Rostral; C: caudal; M: Medial; L: Lateral; D: Dorsal; V: Ventral.

### Growth trajectories of the human, macaque, and mouse corpus callosum

Comparative analyses of growth trajectories can pinpoint what developmental programs might account for variation in cortical association pathways in humans versus mice. To quantitatively investigate whether the corpus callosum growth follows a similar trajectory across primates and rodents, we measured the area of the corpus callosum from FA images of mice, macaques, and humans at several stages of development. We here select FA images from diffusion MR scans, rather than structural MR scans, as we had done when comparing the corpus callosum of adult primates with those of rodents. We select FA scans to track the development of the corpus callosum because the corpus callosum is easily identifiable from its bright white appearance from mid-sagittal sections at successive stages of development and across species. The FA images range from E17 to post-natal day 60 in mice (n=42), from GW 30 to 18 years of age in humans (n=22), and from 2 post-natal weeks to 36 months after birth in macaques (n=102). In humans, we supplement FA images with MR templates ranging from birth to two years of age (Shi et al., 2011), because of the paucity of human MR scans available from 0 to 2 years of age. We supplement these data with a previously published dataset that relied on structural MRI scans to capture the growth of the corpus callosum in humans (n=86) whose ages range from shortly after birth to 25 years of age (Sakai et al., 2017). The FA images of mice are made available by John Hopkins University and the human brain scans were previously published (Chuang et al., 2011). The FA images of macaques are from the UNC-Wisconsin Neurodevelopment Rhesus Database (Young et al., 2017). The FA images of developing human brains were from Boston Children’s Hospital. Details of MR acquisition have been described previously, and these participants were awake when scanned (Cohen et al., 2016). We measured the area of the corpus callosum in mice, rhesus macaques, and humans from mid-sagittal sections at different ages. We excluded one outlier for macaques because the corpus callosum measurement from FA images was inconsistent with that obtained from the T1 structural MR.

We assess whether the corpus callosum growth is extended in primates compared with rodents after controlling for variation in developmental schedules. The timing of developmental transformations (i.e., events) can be used to identify corresponding ages across species (Clancy et al., 2001). Controlling for variation in developmental schedules permits identifying whether certain development processes occur for an unusually long or short period of time in humans (Finlay and Darlington, 1995; Clancy et al., 2001; Clancy et al., 2007; Nagarajan and Clancy, 2008; Workman et al., 2013; Charvet, Hof, et al., 2017; Charvet and Finlay, 2018). Because the translating time resource extends up to 2 years of age in humans and its equivalent in other species, we extrapolated ages at later time points as we had done previously (Charvet et al., 2019). After controlling for variation in developmental schedules between species, we assess whether the corpus callosum ceases to grow later than expected in humans and macaques relative to mice. Statistical analyses used to compare the growth of the corpus callosum between species were performed with the software programming language R. We tested whether a linear plateau would fit the data in each species with the library package easynls (model=3; linear plateau).

## Results

We describe the overall organization of pathways in the forebrain of humans and mice during development with a particular focus on radial and emerging commissural fibers. We also observe that diffusion MR tractography identifies pioneer axons originating from the medial wall of the developing cerebral cortex in both humans and in mice.

### Radial glia organization of the cortex in humans and mice

We explore the development of fiber organization in developing mouse brains from E10.5 to PW 4 (**Fig. 1, Supplementary Fig. 2**) and in humans from GW 17-30 (**Fig. 2–3**). In mice, many of the fibers are observed coursing across the medial to lateral direction in the forebrain throughout development (**Fig. 1, 2**). According to the translating time model, a mouse at E16.5 is roughly equivalent to humans at GW 12-16, and a mouse at E17.5 is equivalent to humans at GW14-18 (**Fig. 2**). We therefore compare mice at E16.5-17.5 to humans at GW 17-21. At GW 17-21, fibers are principally oriented radially with fibers extending from the proliferative pool towards the CP (**Fig. 2A-C**). These fibers likely represent radial glial cells known to guide the migration of neurons from the proliferative pool towards the CP. At E16.5 (**Fig. 2D-E**) as in E17.5 (**Fig. 2F**), we observe fibers coursing principally across the medial to lateral direction through the cortex. Some of these fibers likely correspond to thalamo-cortical, cortico-thalamic or callosally projecting pathways. Thus, diffusion MR imaging highlights that mice and humans matched for age differ in their cortical organization. Radially-organized fibers are evident in the human cortex but this is not the case in mice, presumably because radial fibers dissipate earlier in mice than in humans. Indeed, cortical neurogenesis wanes around E18 in mice (Caviness et al., 2003, 2009), but cortical neurogenesis is thought to persist beyond GW 20 in humans (Zecevic et al., 2005; Malik et al., 2013; Charvet, Hof, et al., 2017).

**Figure 2.**
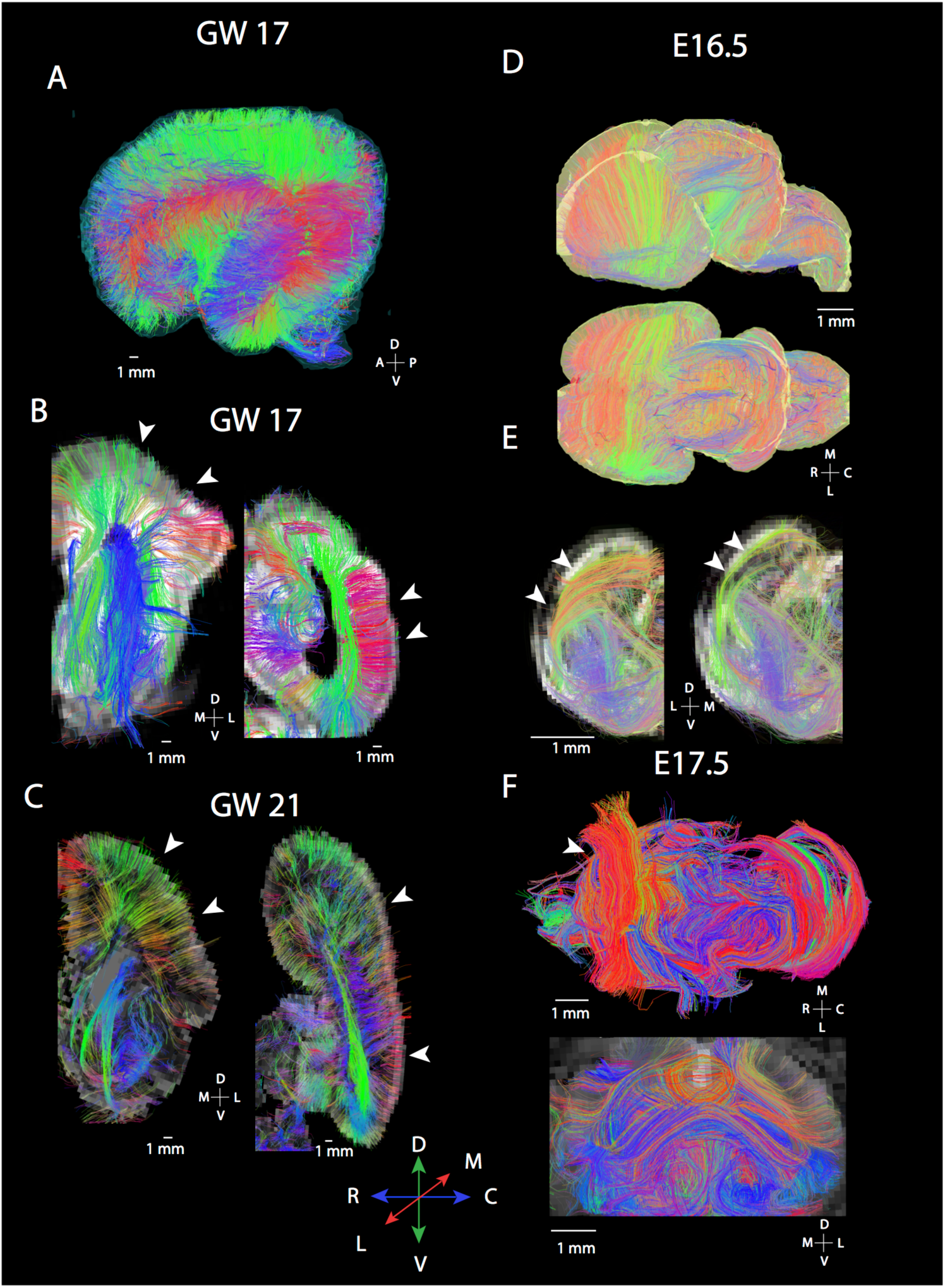
Diffusion MR tractography of cortices at GW17-21 (A-D) and mice (E16.5-E17.6) show major differences in the organization of fibers spanning the cortex. In humans, many fibers course radially within the developing cortex. At GW17 (A-B), coronal slices through the anterior and posterior regions of the cortex show fibers coursing from the proliferative zones to the cortical plates (white arrowheads). A similar situation is observed at GW 21 (C) where coronal slices set to capture fibers identify radial fibers spanning the proliferative pool and the cortical plate in the anterior and posterior cortex (white arrowheads). In contrast, diffusion MR scans of mice at E16.5 (D-E) and at E17.5 (F) show fibers principally organized across the medial to lateral axis (white arrowheads). (E) Coronal slices through the rostral and caudal cortex at E16.5 show fibers coursing tangential to the cortical surface rather than radially (white arrowheads). (F) At E17.5, a similar situation is observed in the mouse where fibers are observed coursing across the medial to lateral direction (white arrowheads). Humans at GW 17-21 and mice at E16.5-17.5 were selected for comparison because they are at roughly equivalent ages of development (Workman et al., 2013). The following abbreviations are used: A: Anterior; P: Posterior; R: Rostral; C: Caudal; M: Medial, L: Lateral; D: Dorsal; V: Ventral.

### Commissural development in mice and humans

In mice, at the earliest gestational age studied (E10.5), pathways crossing the midline were observed in in anterior and ventral regions of the brain **(Fig. 1A-D**). We did not observe callosal fibers at this age. At E14.5 (**Fig. 1E-H**), we observed short pathways crossing the midline in the presumptive corpus callosum (**Fig. 1G-H**). These fibers emerge or terminate within the cingulate cortex, and project contra-laterally through the dorsal midline of the telencephalon (i.e., the developing corpus callosum). We consider these fibers emerging or terminating within the cingulate cortex to represent pioneer axons coursing through the corpus callosum because they are reminiscent of previously described pioneer axons that act as scaffolds to guide development of other axons crossing the dorsal midline (Koester and O’Leary, 1994, Rash and Richards, 2001). At E16.5, fibers course across the medial to lateral direction throughout the developing cortex (**Fig. 1I-J**). We also observe a population of anteriorly located callosal pathways extending across the midline (**Fig. 1K-L**, yellow arrows). Some of these fibers originate or terminate within the cingulate cortex (**Fig. 1K-L**, blue arrows), whereas other fibers originate or terminate from more lateral regions of the developing cerebral cortex. Thus, fibers reminiscent of pioneer axons are first observed emerging or terminating from the cingulate cortex, followed by fibers spanning more lateral regions of the cortex. Given this developmental sequence, neurons emerging from the cingulate cortex may act as a scaffold for later developing neurons that originate from lateral regions of the cortex. As development progresses, the corpus callosum becomes increasingly evident at E17.5 (**Fig. 1M-P**) and PW4 (**Fig. 1Q-T**).

We next compare corpus callosum development in mice (**Fig. 3A-D**) and humans (**Fig. 3E-J**) matched for age. In mice, the corpus callosum is evident from at E16.5 and E17.5 with pathways extending across the medial to lateral axis (**Fig. 3A-D**). At E17.5, a larger callosal structure is evident in the rostral cortex compared with the caudal cortex (**Fig. 3D-C**). In humans, at GW17 (**Fig. 3E**, pink arrow), at GW20 (**Fig. 3G**) through GW22 (**Fig. 3H**) and GW 30 (**Fig. 3I**) we observe some pathways extending from the superior medial wall of the developing cortex projecting towards the contralateral hemisphere through the corpus callosum. Interestingly, other fibers arising from more lateral regions of the cortex also project towards the corpus callosum at GW 17 (**Fig. 3E**, yellow arrow), at GW21-22 (**Fig. 3H**, green arrow), and at GW30 (**Fig. 3I**, white arrow). Some of these fibers course within the subventricular and ventricular zones (**Fig. 3E**, blue arrow). A few fibers extend into the grey matter of the medial wall of the developing cortex at GW 17 (**Fig. 3E**, red arrows), GW 21 and GW 22 (**Fig. 3H**, orange arrows). We also observe variation in corpus callosal structure across the anterior to posterior axis with seemingly larger callosal structures located in the frontal cortex than in more posterior regions of the cortex at GW 17 (**Fig. 3E**). This variation in corpus callosum structure across the anterior to posterior axis of the cortex observed in the GW 17 human is reminiscent of those observed in mice at E17.5. Thus, in humans as in mice, there is a tendency for more anterior callosal pathways to be expanded than in posterior regions. As development progresses, the corpus callosum increasingly extends across the medial to lateral axis in both species.

**Figure 3.**
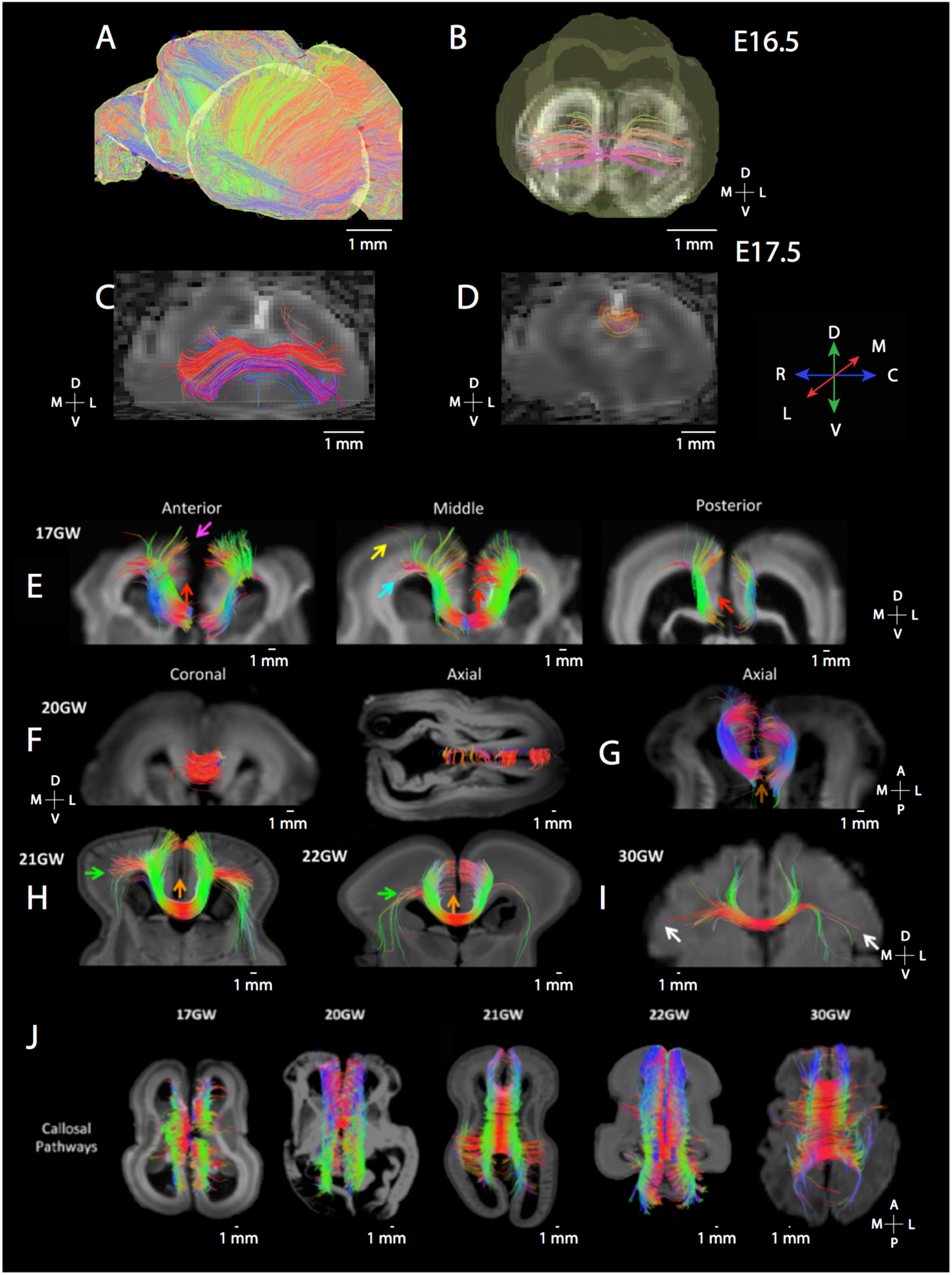
Diffusion MR tractography captures the maturation of the corpus callosum in mice (E16.5:A-B; E17.5:C-D) as well as human brain from GW 17 to 30 (E-J). (A) Sagittal slices set through the E16.5 mouse brain show that fibers are principally organized across the medial to lateral axis. (B) ROIs set through the dorsal midline highlight callosal fibers at this age at E16.5 (B) and at E17.5 (C). The corpus callosum is more prominent at E17.5 than at E16.5. At E17.5, more fibers are evident in the rostral cortex (C) than in more caudal regions (D). At GW17 in humans (E), putative pioneer and callosal pathways are also observed across the anterior to posterior axis with more prominent collosal fibers evident in anterior regions than in posterior regions. (E) The yellow arrow highlights callosal pathways from the lateral brain regions, and the pink arrow shows callosal pathways from the upper medial wall (E) The blue arrow shows callosal pathways from the ventricular region, and red arrows show possible residual pioneer axons at the cingulate cortex. The corpus callosum gains increasing prominence with age (F-I). (H) At GW21, callosal pathways are evident and from the upper medial wall and lateral brain regions (green arrow). (H) Callosal pathways from the upper medial wall, cingulate cortex, and lateral brain regions (green arrow) at GW21 and GW22. (F) At GW 30, we observe callosal pathways from the cingulate cortex, upper medial walls, and lateral walls of the brain (white arrows). (J) Dorsal views of horizontal slices through the corpus callosum from GW17 to GW30 show that the corpus callosum becomes increasingly prominent with age.

### Corpus callosal growth trajectories across species

To test whether there are deviations in the timing of corpus callosum development between humans and mice, we measured the area of the corpus callosum at successive ages across primates (i.e., humans, macaques) and mice. Ages in humans and macaques are mapped onto mouse age to control for variation in developmental schedules across species. We use the translating time model to find corresponding ages, which relies on the timing of developmental transformations to find corresponding ages across species (Clancy et al., 2001; Workman et al., 2013; Charvet, Hof, et al., 2017; Charvet and Finlay, 2018).

In both humans and in mice, the corpus callosum grows and ceases to grow (i.e., plateaus) after birth (**Fig. 4**). In humans, the corpus callosum grows and reaches a plateau at about 1.64 years of age in humans, which is equivalent to 53.5 days after conception in mice (easynls fit model=3; adj R^2^=0.78; p<0.05; n=22; **Fig. 4A**). In macaques, the corpus callosum continues to grow past the predicted ages for those of mice (post-natal day 12.5) but our non-linear model does not capture a linear plateau with these data (easynls fit model=3; n=102; **Fig. 4B**). In mice, the corpus callosum growth ends at around 31 days after conception, which is 12.5 days after birth (easynls fit model=3 adj R^2^=0.81; p<0.05; n=42; **Fig. 4C**). Because of the small sample available for study between the ages of birth to 2 years of age in humans, we supplement our analysis with a previously published dataset that captures the growth of the corpus callosum in humans to ensure the validity of these results (Sakai et al., 2017; **Supplementary Fig. 3**). Such an analysis shows that the corpus callosum growth plateaus at 1.87 years of age (easynls fit model=3; adj R^2^=0.78; p<0.05; n=86; **Supplementary Fig. 3**). These two different datasets both demonstrate that corpus callosum growth ceases around 1-2 years after birth in humans. That is, the corpus callosum continues to grow for an extended period of time in humans compared with mice.

**Figure 4.**
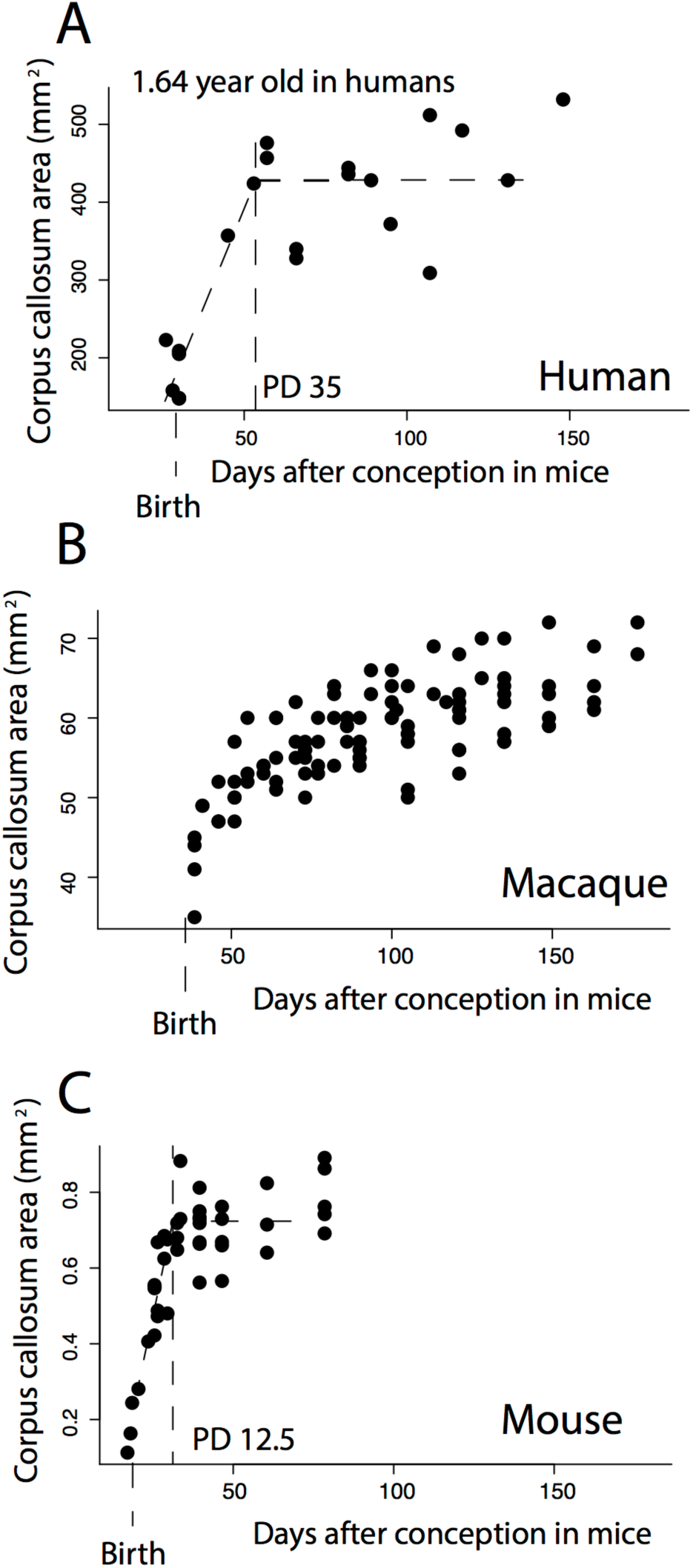
The growth of the corpus callosum is protracted in human (A), and macaque (B) compared with mice (C). In order to control for variation in the length of developmental schedules, age in humans and macaques is mapped onto mouse age in days after conception according to the translating time website (Clancy et al., 2001; Workman et al., 2013). (A) After controlling for variation in the length of development across species, the area of the corpus callosum reaches a plateau between 1 to 2 years of age in humans (i.e., 1.64 years after birth). (B) The corpus callosum shows sustained growth in macaques across the examined ages and does not plateau. (C) The corpus callosum reaches a plateau at 31 days after conception in mice, which is 12.5 days after birth in mice. The corpus callosum grows for much longer than expected in humans and macaques when human and macaque age is mapped onto that of mice. Vertical bars identifies when the corpus callosum ceases to grow, as well as birth.

We next confirm the protracted growth of the corpus callosum statistically, and we select the most recent and largest dataset capturing developmental transformations across rodents, and primates, including humans (Charvet, Šimić, et al., 2017). Neural transformations that occur on 30-32 days after conception in mice (n=10), which is when the corpus callosum ceases to grow in mice, occur between GW 20 to about 3.5 months after birth in humans (Lower 95%CI: GW 21; Upper 95CI: 2.7 months after birth). Yet, in humans, the corpus callosum growth ends at 1.64 years of age, which is past the 95% confidence intervals generated from these data. Taken together, these findings demonstrate that the corpus callosum continues to grow for significantly longer than expected in humans compared with mice.

## Discussion

In this study, we tracked the development of fibers in the developing human and mouse cortex. Our findings demonstrate the ability of HARDI to track fiber development in different species. During cortical neurogenesis, fibers of the developing human cortex are primarily radially aligned but that this is not the case in age-matched mice. We focus on the developmental timeline of corpus callosum development, and we find that the corpus callosum grows for an extended period of time in humans compared with mice. We discuss the strengths and limitations of diffusion MR tractography, as well as the potential implications for the observed conserved and deviant developmental programs giving rise to variation in cortical structure and connectivity patterns in the human lineage.

### Diffusion MR scanning for basic development and health

Although ultrasound remains the predominant modality for evaluating disorders related to pregnancy, fetal MRI has been increasingly used to track normal and abnormal human development (Mitter et al., 2015; Jakab et al., 2017). Diffusion MR imaging is still in its infancy, and more work is needed to identify what developmental processes can be observed from diffusion MR tractography. HARDI is a new innovative approach to identify developmental programs in the human fetal brain, which can also be used to identify conserved and deviant developmental programs between humans and model organisms such as mice. It is important, however, to acknowledge the reliability of HARDI tractography when reporting diffusion imaging studies. As this method is in its infancy, HARDI does suffer from certain pitfalls (Maier-Hein et al. 2016). According to Maier-Hein et al., currently available algorithms for fiber pathway reconstruction from *in vivo* scans tend to produce a number of false positives. While our algorithms are similar to the ones used by Maier-Hein et al., we reason that the architecture of the fetal telencephalon, shorter fiber pathways, and probably fewer total fiber crossings during ongoing axonal elongation, lead to a significantly smaller number of reconstructed false positive pathways from *ex vivo* scans. Additionally, the low number of specimens available for this study makes it challenging to draw quantifiable conclusions from HARDI tractography. It is currently difficult to obtain sufficient postmortem fetal brains to resolve this problem. Due to these limitations, we integrate our data with comparative analyses of callosal growth trajectories. However, studies such as these are essential as they identify key developmental processes (e.g., pioneer axons) in the human fetal brain, which should increase the utility of this work for disease detection and prevention.

### Development of radial organization in humans and mice

Radial glia have been identified as an important player in developmental mechanisms accounting for the expanded isocortex of primates in adulthood (Rakic, 1978,1995), and we observe major differences in radial organization across the isocortex of humans and mice at corresponding ages. Fibers coursing across the mouse cortex from E16.5-17.5 are principally organized across the medial to lateral axis. These fibers, which extend across the medial to lateral axes of the developing cortex, likely represent cortico-thalamic or thalamo-cortical pathways. This is in contrast with humans at corresponding ages where fibers in humans (GW17-20) are primarily organized radially across the isocortex. The radial organization of the developing cortex has previously been characterized with diffusion MR tractography in humans. Radially-aligned fibers are evident at GW 17, and begin to diminish around GW 21 with residual fiber pathways still observed at GW 40 (Miyazaki et al., 2016). As development progresses, radial glia as well as other proliferative cells wane as cells exit the proliferative zone, and newly born neurons assume their final position within the cortex (Rakic, 1974; Miyata et al., 2001; Zecevic et al., 2005; Takahashi et al., 2012; Malik et al., 2013; Arshad et al., 2015; Charvet, Hof, et al., 2017).

The finding that the human fetal cortex shows a predominance of fibers oriented radially compared with that observed in mice can be integrated with known deviations in developmental programs between the two species. After controlling for variation in developmental schedules, humans and macaques extend the duration of cortical neurogenesis. It is not the onset of cortical neurogenesis that is delayed in primates relative to rodents, but rather, it is the end of cortical neurogenesis that is protracted in primates relative to rodents (Clancy et al., 2001; Cahalane et al., 2014). Extending the duration of neuron production should lead to an extension in the duration in which neurons are produced and migrate to the cortical plate. Accordingly, mice at E16.5 and E17.5 should not possess many radially aligned fibers because neurogenesis is largely complete at these ages (Caviness et al., 2003, 2009; Yuzwa et al., 2017; Preissl et al., 2018). This is in contrast with humans at GW17-21, which still possess a large proliferative pool, self-renewing radial glia, as well neurons exiting the proliferative pool to migrate towards the cortical plate (Rakic, 1974, 2003; Zecevic et al., 2005; Malik et al., 2013; Arshad et al., 2015; Charvet, Šimić, et al., 2017). Thus, the observation that radial glia are evident in humans but not in age-matched mice aligns with known deviations in the duration of cortical neuron production between primates and rodents.

### Development of commissural fibers in mice and humans

We track the developmental sequence of corpus callosum development in mice. We observed fibers reminiscent of pioneer neurons emerging or terminating from the cingulate cortex as early as E14.5. Similar fibers were also observed at E16.5 (Koester and O’Leary, 1994, Molnár et al., 1988; McConnell et al., 1989; Ozaki and Wahlsten, 1992; 1998; Koester and O’Leary, 1994; Rash and Richards, 2001; **Fig. 1C**). Although it is now well known that callosally-projecting pioneers emerge from the cingulate cortex, the origin of pioneer neurons crossing through the dorsal midline of the telencephalon (i.e., the presumptive corpus callosum) was the subject of debate in the 90s. The debate focused on whether pioneer neurons crossing the dorsal midline originate from the cingulate cortex or from more lateral regions of the cortex. The debate was settled with the use of dies, which showed that pioneer neurons have their cell bodies originate in the cingulate cortex rather than more lateral regions of the cortex (Rash and Richards, 2001). Since then, more work has tracked the development of pioneer neurons extending axons to the contralateral hemisphere (Chun and Shatz, 1989; McConnell et al., 1989; Chun et al., 1987; Antonini and Shatz, 1990; Imai and Sakano, 2011).

In humans as in mice, we identify putative pioneer neurons during development, which can be imaged as early as GW 17. These data are in line with the work of others who have shown that neurons emerging from the developing cingulate cortex have axons crossing the dorsal midline, which can be identified immunohistochemically in the human fetal brain at these ages (Ren et al., 2006). At GW 17, short fibers originating from the upper medial wall of the developing cortex and, more sparsely, from the cingulate cortex course towards the contralateral hemisphere in humans (**Fig. 3E**, see pink arrow). The sparse fibers from the cingulate cortex likely correspond to residual pioneer pathways, which are known to originate in the cingulate cortex and cross the midline (Rakic and Yakovlev, 1968; Koester and O’Leary, 1994, Ren et al., 2006). As development progresses, the corpus callosum becomes increasingly evident in both humans and mice.

### Conservation and variation in the timing of corpus callosum development

We compare developmental timing of the corpus callosum in mice, macaques, and humans (Schwartz and Goldman-Rakic, 1991; Clancy et al., 2017; Workman et al., 2013). According to the translating time model, early callosal developmental milestones such as the emergence of the corpus callosum, and myelination onset are conserved in their timing across humans and mice (Workman et al., 2013). In the present study, we do not observe major differences in callosal development between the two species at relatively early stages of its development, although these findings do not necessarily entail there are no early differences across species. We do, however, find that there are clear differences in the growth trajectories of the corpus callosum between primates and mice such that deviations in the duration of callosal growth account for the expansion of the corpus callosum in primates.

After controlling for variation in overall developmental schedules across species, the corpus callosum continues to grow for an extended period of time in humans. We find that the corpus callosum ceases to grow somewhere between 1 to 2 years of age, which is consistent with the work of others who have shown that the corpus callosum ceases to grow several months to a few years after birth in humans (Giedd et al., 1999; Tanaka-Arakawa et al., 2015; Vannucci et al., 2017; Sakai et al., 2017). In mice, the corpus callsoum area only grows for a few days, which is also consistent with previous work (Wahlsten, 1984; but see Chuang et al., 2011). We found that the macaque corpus callosum grows throughout the examined postnatal ages, and these findings align with previous work tracking the growth of the corpus callosum in macaques (Scott et al., 2016). We have not here identified the underlying developmental processes responsible for variation in callosal growth trajectories between species and several possibilities exist. The human corpus callosum might prolong the duration with which corpus callosum axons increase in number, or axons increase in thickness. Alternatively or in addition, the duration with which axons myelinate may be protracted in humans relative to mice. Although we do not resolve the underlying cellular process responsible for variation in corpus callosal growth trajectories across species, our findings demonstrate that the protracted development of cortico-cortical pathways are linked to their expansion in the primate lineage (Charvet, Šimić, et al., 2017, Charvet et al., 2019).

The corpus callosum is responsible for a variety of complex behaviors, and protracted growth of long-range cortico-cortical pathways such as the corpus callosum may be responsible for protracted behavioral development observed in humans relative to mice. Some behaviors such as weaning are deviant in their timing in humans relative to other species (Hawkes and Finlay, 2018). However, very studies have compared the timing of behavioral development between humans and other species. Integrating data on the timing of long-range cortico-cortical pathways with behavioral milestones may serve to identify the behavioral implications of extending the maturation of long-range cortico-cortical pathways in the primate lineage.

The timing of developmental transformations may be a powerful mechanism through which to generate variation in cortical association pathways. Several hypotheses have focused on the timing of axonal development as a source of evolutionary changes in connectivity patterns (Innocenti, 1995; Striedter, 2005). It has been proposed that neurons that reach their targets early can outcompete other targets, thereby expanding their representation within the brain (Catania, 2001). A previous study showed that the star-nose mole’s fovea matures earlier than adjacent appendages. Thus, early axon extension might favor competing for targets, and their over-representation in the mature isocortex (Catania, 2001). On the other hand, prolonging the duration of axon extension and refinement might afford more time to outcompete targets, which would accordingly increase territory of select pathways in the adult brain (Deacon, 1990; Striedter, 2005). These two seemingly contradictory hypotheses emphasize the importance of timing in generating variation in connections between species. Our study supports the view that an extended duration with which corpus callosal pathways grow account for evolutionary changes in their structure in the adult brain. Thus, evolutionary changes in a cortical association pathway emerge by modifying the time with which they develop.

## Acknowledgements

This work was supported by NICHD (R01HD078561, R21HD098606, R21HD069001; ET), NIMH (R21MH118739; ET) and NINDS (R03NS091587; ET). This research was carried out in part at the Athinoula A. Martinos Center for Biomedical Imaging at the Massachusetts General Hospital, using resources provided by the Center for Functional Neuroimaging Technologies, NIH P41RR14075, a P41 Regional Resource supported by the Biomedical Technology Program of the National Center for Research Resources (NCRR), National Institutes of Health. This work also involved the use of instrumentation supported by the NCRR Shared Instrumentation Grant Program (NIH S10RR023401, S10RR019307, and S10RR023043) and High-End Instrumentation Grant Program (NIH S10RR016811). Some of this work was also funded by an NIGMS grant (5P20GM103653) for research at Delaware State University. A developmental series of mouse FA scans were obtained as a courtesy of Dr. Mori (lbam.med.jhmi) at Johns Hopkins University. Dr. Lana Vasung provided comments on a previous version of this manuscript. This study was conducted partly using postmortem human brain specimens from the tissue collection at the Department of Neurobiology at Yale University School of Medicine (supported by grant NIH MH081896), which form a part of the BrainSpan Consortium collection (http://www.brainspan.org). Dr. Rebecca D. Folkerth at Brigham and Women’s Hospital provided the other brain specimens.

