## Supplementary Fig. 1 for "High angular resolution diffusion MRI reveals conserved and deviant programs in the paths that guide human cortical circuitry"

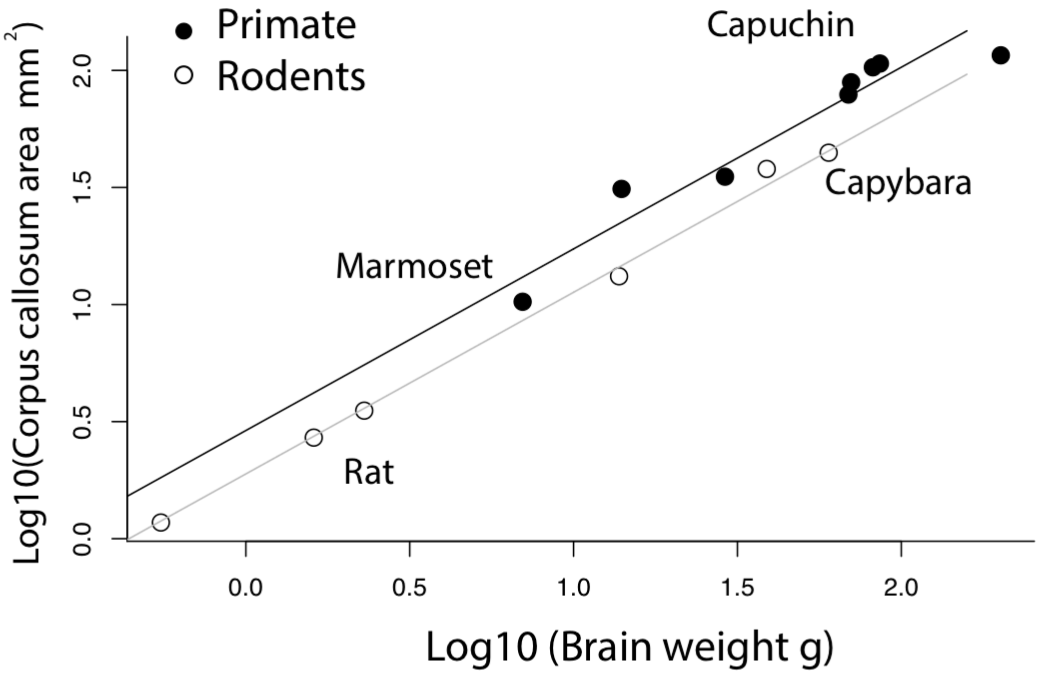


**Supplementary Figure 1.** For a given brain size, the corpus callosum area is relatively expanded in primates compared with rodents. Phylogenetically-controlled regressions are obtained separately for primates and rodents.
