## Supplementary Fig. 2 for "High angular resolution diffusion MRI reveals conserved and deviant programs in the paths that guide human cortical circuitry"

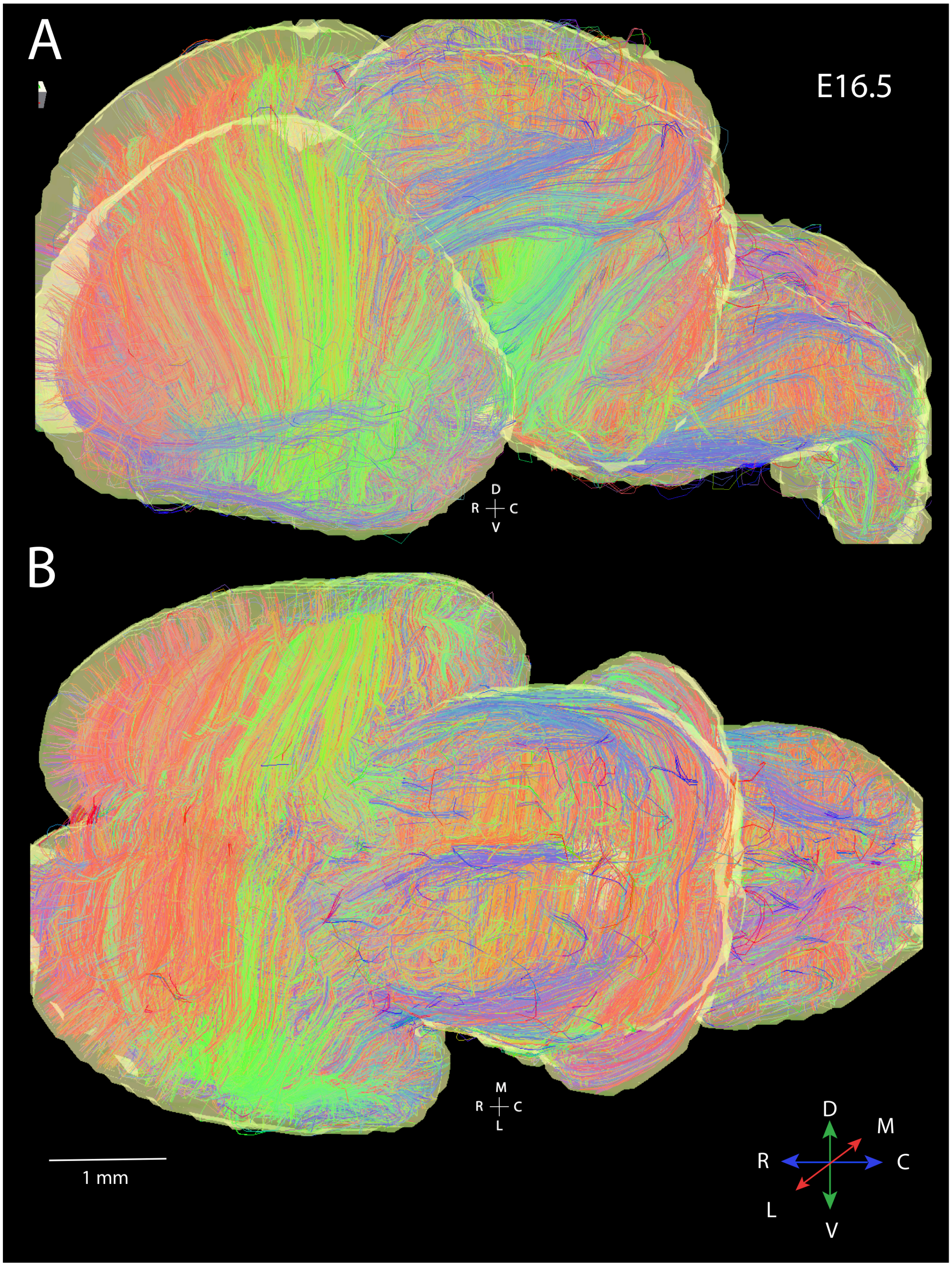


**Supplementary Figure 2.** Whole brain MR tractography at E16.5 demonstrates that fibers are course primarily across the medial to lateral direction within the developing cortex. Minimum length threshold of fibers was set to 0.8mm. R: Rostral; C: caudal; D: Dorsal; V: Ventral; M: Medial; C: Caudal.
