## Supplementary Fig. 3 for "High angular resolution diffusion MRI reveals conserved and deviant programs in the paths that guide human cortical circuitry"

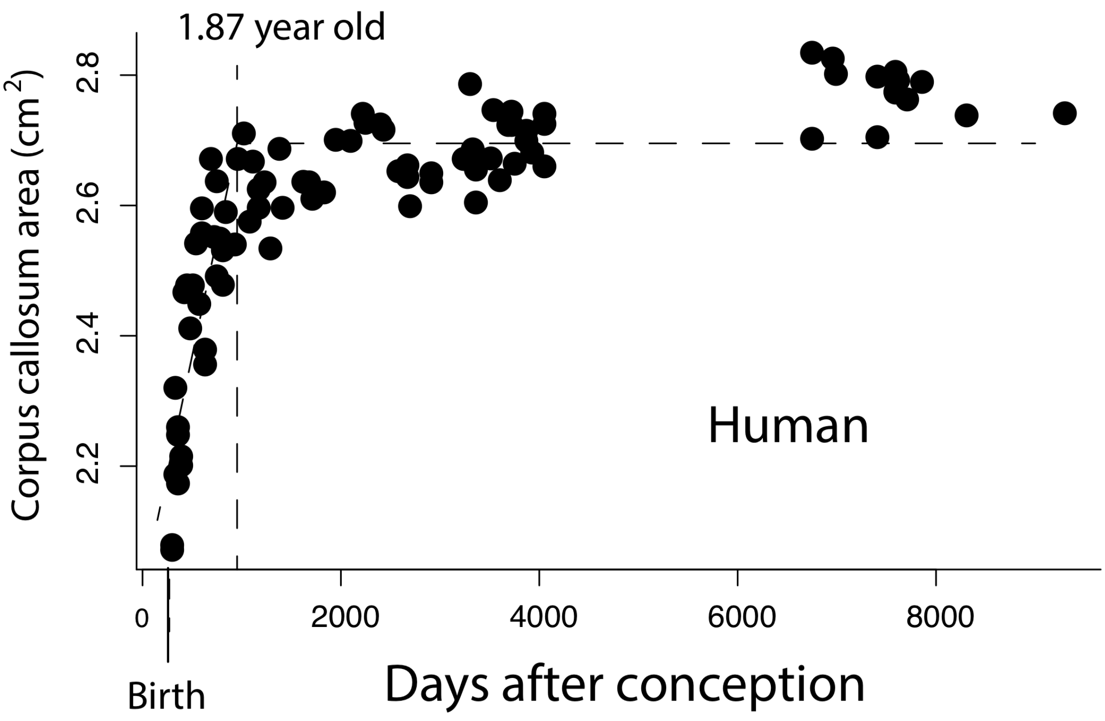


**Supplementary Figure 3.** The corpus callosum area grows after birth to reach a plateau between 1 to 2 years of age (i.e., 187 years old), which is consistent with those reported in the present study. These data are from structural MRI scans and are from Sakai et al., 2017. Vertical bar identifies when the corpus callosum ceases to grow, as well as the time of birth.
